# Interferon-α and -β subtypes have temporally distinct roles in containing viral spread and protecting vital organs

**DOI:** 10.64898/2026.03.09.710658

**Authors:** Carolina R. Melo-Silva, Marisa I. Roman, Natasha Heath, Michelle Gelman, Megan Keller, Jihae Choi, Lingjuan Tang, Oluwatomilola Taiwo, Samita Kafle, Edilson Torres-González, Holly Ramage, Raul Andino, Luis J. Sigal

**Affiliations:** Department of Microbiology and Immunology, Thomas Jefferson University, Philadelphia, PA 19107, USA; Department of Physics, St Joseph University, Philadelphia PA 19131, USA; GlaxoSmithKline, 1250 S. Collegeville Road, Collegeville, PA 19426, USA; Merck&Co Inc, 770 Sumneytown Pike, West Point PA 19486, USA; Department of Microbiology and Immunology, University of California San Francisco, San Francisco, CA 94158, USA

**Author notes:** Carolina R Melo-Silva, Marisa I Roman, Natasha Heath, Michelle Gelman, Megan Keller, Jihae Choi, Lingjuan Tang, Oluwatomilola Taiwo, Samita Kafle, Edilson Torres-González, Holly Ramage, Raul Andino.

## Abstract

Type I interferons (IFN-Is), including a single IFN-β and over a dozen IFN-αs, induce the anti-viral state. *In vitro*, IFN-β transcription requires the assembly of the “enhanceosome” composed of the constitutive transcription factors NF-κB and AP-1, and either constitutive IRF3 or IFN-I-inducible IRF7. IRF3 and IRF7 transcribe mouse IFN-α4 and human IFN-α1/13. Only IRF7 transcribes other IFN-αs. How IFN-I subtype multiplicity and their differential constitutive/IFN-I-inducible versus only IFN-I-inducible transcription help control viruses *in vivo* remains unknown. Using novel genetically modified mice, we demonstrate that most or all IFN-I subtypes, regardless of their transcriptional control, are necessary to curb the systemic dissemination of lymph-borne ectromelia virus (ECTV) but do not necessarily suppress ECTV or West Nile Virus (WNV) replication in the liver or the brain, or promote survival to their infection. Individually, the most critical IFN-I subtype to survive ECTV and WNV infections is IFN-β. IFN-α4 potentiates IFN-β but is not essential. PRDII, the IFN-β promoter’s NF-κB binding site, which is required for enhanceosome assembly, is dispensable for *in vivo* IFN-β production but complements IRF7-IFN-β transcription to restrain ECTV and WNV. IFN-β alone protects IFN-α-deficient mice from ECTV but not from WNV lethality. Control of WNV replication in the brain successively requires IFN-α, IFN-β, and then PRDII-dependent IFN-β, suggesting that brain protection requires IFN-I production by various cell types and pathways. Thus, contrary to the prevailing view, *in vivo*, IFN-β does not function earlier than IFN-α, and IFN-I subtypes non-redundantly cooperate to restrain viruses in the periphery and in vital organs.

## Introduction

Type I Interferons (IFN-I) are small cytokines encoded within the IFN-I locus by multiple intron-less genes that bind to the common Interferon-Alpha-Receptor (IFNAR) ^1–4^. IFNAR signaling induces the transcription of hundreds of interferon-stimulated genes (ISGs) that help restrict viral replication and activate the immune system ^5,6^.

In primates and rodents, the IFN-I gene complex encodes a single IFN-β and multiple IFN-α (thirteen in humans and fourteen in mice) subtypes. Genetic studies concluded that the paralogous expansion of the IFN-I genes results from gene conversion and duplication, suggesting that, in these orders, there is an independent evolutionary drive to increase the number of IFN-α genes but not the number of IFN-β genes ^7^.

In humans and mice, the IFN-β gene, respectively IFNB1 and *Ifnb1*, is regulated by almost identical Positive Regulatory Domains (PRD), which can be bound by the constitutive transcription factors (TF) NF-κB, AP-1, and IRF3, and the IFN-I inducible IRF7 (**Fig. 1A**) ^8–10^. In cultured cells, AP-1, IRF3/7, and NF-κB must bind simultaneously to the IFNB1/*Ifnb1* promoter together with the high mobility architectural protein HMG I(Y), to form an “enhanceosome” that alters chromatin accessibility for IFN-β production ^9–13^. The IFN-α genes, IFNA in humans and *Ifna* in mice, are regulated by promoters containing up to four Virus Responsive Elements (VRE-A, -B, -C, and -D). VRE-C is the only VRE that can be bound by either constitutive IRF3 or IFN-I-inducible IRF7 and is only present in the promoters of *Ifna4* in mice and of IFNA1 and IFNA13 in humans (**Fig. 1B**) ^8,14–16^. VRE-A, -B, and -D are only bound by IFN-I-inducible IRF7. Consistent with IRF7 being an ISG in most cells, all *Ifna* genes, except *Ifna4*, require positive feedback for their transcription in mouse fibroblasts ^17^. Thus, based on their transcriptional regulation, we can distinguish: **1)** IFN-I subtypes transcribed by **c**onstitutively- and **I**FN-I-**i**nducible TFs (C&II IFN-I subtypes), which are traditionally known as “early” IFN-I subtypes. In mice, these are IFN-β and IFN-α4 (IFN-β and IFN-α1/13 in humans); and **2)** IFN-I subtypes transcribed only by **I**FN-I-**i**nducible IRF7 (II IFN-I subtypes), traditionally known as “late” IFN-I subtypes because their transcription requires de novo IRF7 synthesis. In mice, these are all the IFN-α subtypes except for IFN-α4.

**Figure 1.**
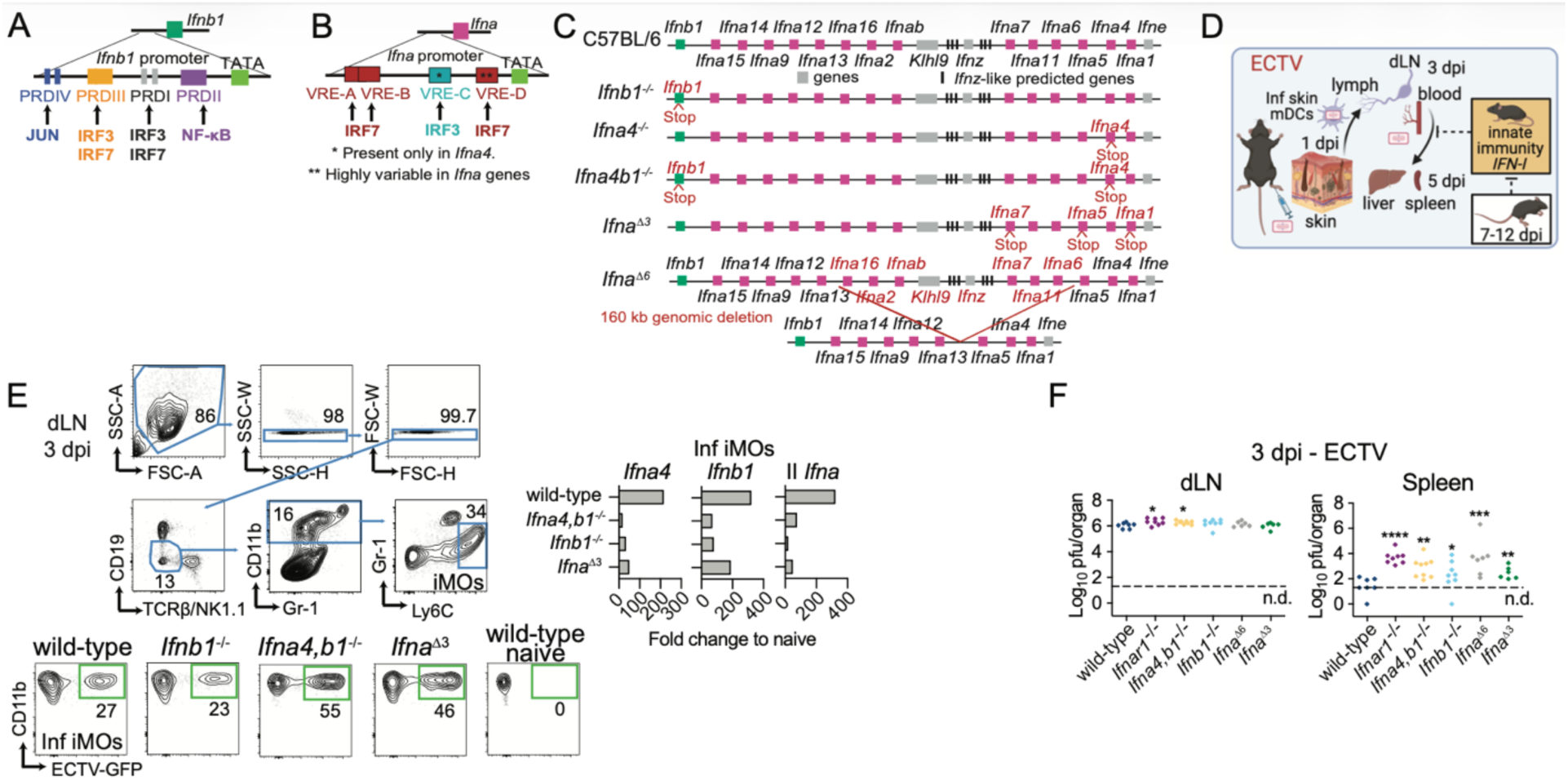
C&II and II IFN-I subtypes are required to restrict ECTV systemic dissemination and to sustain positive feedback for IFN-I production *in vivo*. **A-B**) Transcriptional regulation of mouse *Ifna* (**A**) and *Ifnb1* (**B**) genes, described in the main text. **C**) Genetic map diagrams of the IFN-I gene complex in C57BL/6, *Ifnb1^−/−^*, *Ifna4^−/−^*, *Ifna4b1*^−/−^, *Ifna^Δ3^*, and *Ifna^Δ6^* mice made by CRISPR and iGONAD ^27^. **D**) ECTV intra-host dissemination after subcutaneous footpad inoculation ^29^ (created with BioRender). **E**) IFN-I transcription quantified by RT-qPCR of RNA extracted from FACS-sorted infected iMOs (GFP^+^) at three dpi from pooled popliteal dLN of the indicated mice infected in both rear footpads with 3,000 pfu of ECTV-GFP. The data is representative of three independent experiments with similar outcomes in which each IFN-I-deficient strain was tested at least twice. **F**) ECTV titers in the popliteal dLNs and spleens quantified by plaque assay from the indicated mice at three dpi following footpad infection with 3,000 ECTV pfu. The data correspond to two independent experiments combined. Each symbol depicts an individual mouse. P values were calculated compared with wild-type controls using the t-test with Welch’s correction. The dotted line indicates the plaque assay detection limit (n.d.: not detected).

IFNAR is a heterodimeric receptor composed of IFNAR1 and IFNAR2. IFN-β binds IFNAR1 with higher affinity than any IFN-α subtype and through a different amino acid interface ^18–20^. As a result, IFN-α and IFN-β may induce different ISG subsets and differ in some of their anti-viral functions. Moreover, IFN-α subtypes exhibit variable affinities for IFNAR, which may affect their antiviral effects ^18–22^. For example, exogenous treatment with different IFN-α subtypes variably suppresses hepatitis B virus, retroviruses, and SARS-CoV-2 ^23–26^. However, whether IFN-α multiplicity is necessary to control viral infections, or whether individual endogenous IFN-I subtypes and their specific transcriptional regulation play unique or redundant roles *in vivo*, remains unknown. To begin unraveling these questions, we created a collection of IFN-I subtype-deficient mouse strains. We used them to unveil the roles of IFN-I subtypes and their transcriptional regulation in controlling two lymph-borne viral infections.

## Results

### 1. C&II and II IFN-I subtypes are required to restrict ectromelia virus systemic dissemination and to sustain positive feedback for IFN-I production in vivo

To understand the biological role of IFN-I subtypes and their transcriptional regulation, we created genetically modified C57BL/6 (B6, herein wild-type) mice deficient in C&II IFN-β (*Ifnb1*^−/−^), both C&II IFN-I subtypes (*Ifna4,b1*^−/−^), and three II IFN-α genes (*Ifna*^Δ3^, lack IFN-α1, -α5, and -α7) by introducing stop codons. We also made mice deficient in six II IFN-α subtypes (*Ifna^Δ6^*). These mice have a deletion that includes IFN-α16, -α2, -αb, -α7, -α11, and -α6, and IFN-ζ and the ubiquitination adaptor KLHL9 (**Fig. 1C**) ^27^. To mimic a natural infection, we infected mutant and wild-type mice in the footpad with the double-stranded DNA *Orthopoxvirus* ectromelia virus (ECTV), which models human smallpox and mpox ^28,29^. ECTV lymph-borne infection can be temporally divided into a peripheral phase, when infection is restricted to the foot and the popliteal draining lymph node (dLN), and a systemic phase, when the virus invades the spleen and liver through efferent lymphatics and the bloodstream (**Fig. 1D**).

At three days post-infection (dpi), infected inflammatory monocytes (iMOs) are the predominant cells that transcribe various IFN-I subtypes in the dLN of wild-type mice infected with ECTV ^30,31^. At this time point, *Ifnar1^−/−^*iMOs scarcely transcribe C&II and II IFN-I (90-95% reduction), indicating that this localized, early IFN-I transcription requires IFN-I positive feedback ^31,32^. To test the roles of C&II and II IFN-I in this positive feedback, we sorted infected iMOs from wild-type, *Ifna4,b1*^−/−^, *Ifnb1*^−/−^, and *Ifna^Δ3^* at three dpi with ECTV, and compared IFN-I transcription by RT-qPCR. In these mutant strains, the targeted IFN-I genes are not translated but can still be transcribed. The transcription of C&II *Ifnb1*and *Ifna4,* and II *Ifna* was decreased in infected iMOs from all mutant strains compared to wild type mice (**Fig. 1E**). These results reveal that even minor disruptions of the IFN-I locus severely alter iMOs’ early transcription of IFN-I in the dLN, and that, contrary to the prevailing view, not only C&II IFN-β and IFN-α4 but also II IFN-α subtypes provide positive feedback for overall early IFN-I transcription in the dLN.

Wild-type mice are highly resistant to ECTV lethality. IFNAR1 deficiency results in slightly increased ECTV loads in the dLN at two to three dpi, early presence of ECTV in the liver or spleen at three dpi, and full lethality by seven to eight dpi ^32–36^. Thus, we compared virus loads in the dLNs and spleens of wild-type and the IFN-I-deficient strains at three dpi. Compared to wild-type, ECTV titers were slightly higher in the dLNs from *Ifnar1*^−/−^ and *Ifna4,b1*^−/−^ but not from *Ifnb1*^−/−^, *Ifna^Δ6^*, or *Ifna^Δ3^*mice (**Fig. 1F**), indicating a minor but significant role for the combined C&II IFN-I in restricting viral replication in the dLN. Notably, compared to wild-type mice, all mutant strains had higher viral loads in their spleens. Thus, C&II and II IFN-I subtypes maximize IFN-I transcription in iMOs by positive feedback and non-redundantly restrict the dissemination of ECTV from the dLN to the spleen.

### 2. C&II IFN-I subtypes are essential for controlling lymph-borne viral infections after dissemination, with IFN-β playing a critical role in survival

Most *Ifna^Δ3^* mice survived ECTV infection with 3,000 pfu and did not differ from wild-type mice in their survival frequency (**Fig. 2A**). As expected ^30,34,37^, all control ECTV-susceptible Toll-like receptor-9-deficient (*Tlr9^−/−^*) mice rapidly succumbed to the infection. Consistent with the survival data, *Ifna*^Δ3^ and wild-type mice presented similar viral loads, irrespective of sex, at five, seven, and ten dpi in livers and spleen (**Fig. 2B**). Females of both strains had lower ECTV titers at seven dpi than males (**Fig. 2 C-D**). At ten dpi, females had cleared while several males still had detectable ECTV in the livers and spleens. *Ifna^Δ3^* and wild-type mice were also similarly resistant to 30,000 and 300,000 ECTV pfu (**Fig. 2E**). Thus, deficiency in three II IFN-α subtypes does not compromise survival to ECTV.

**Figure 2.**
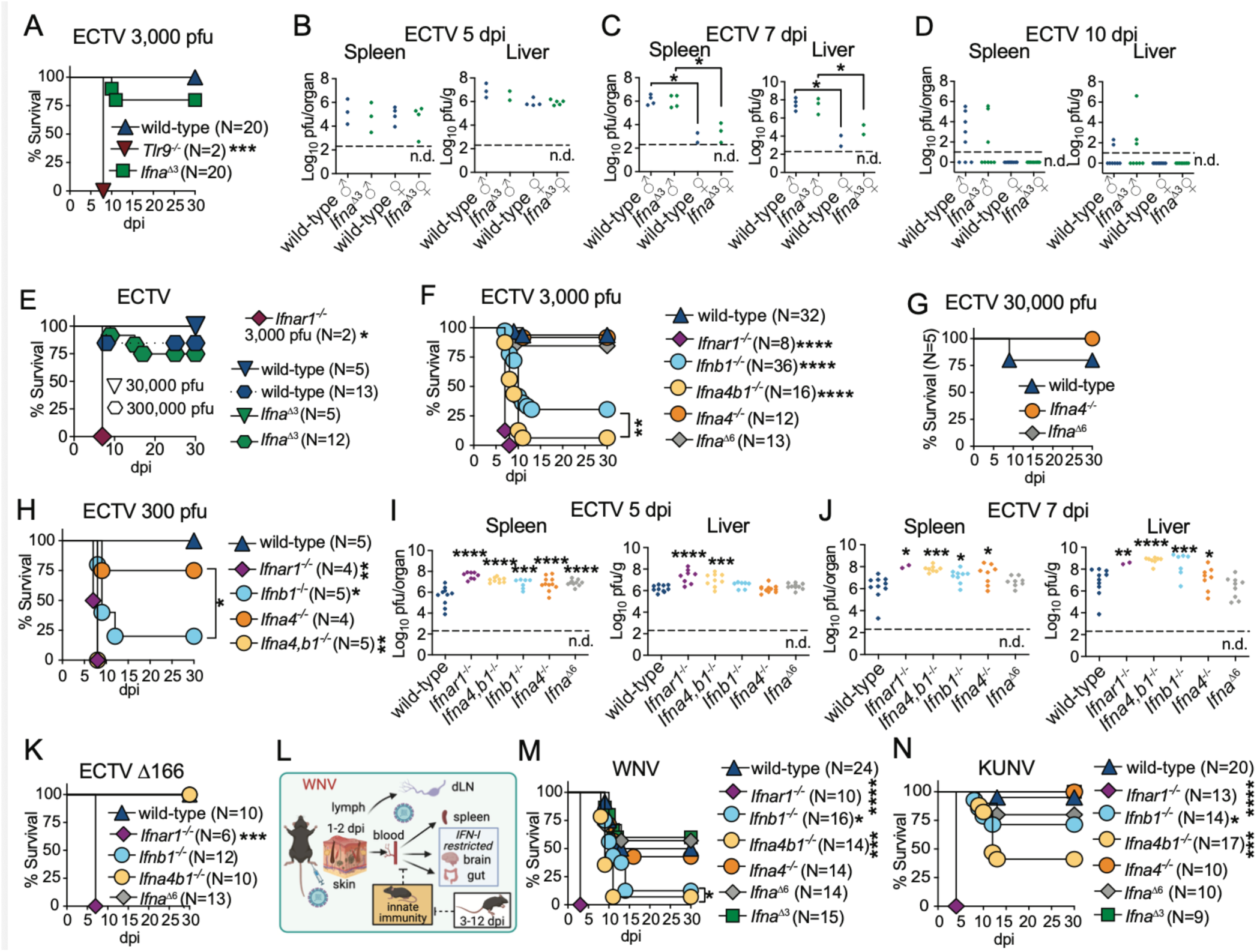
C&II IFN-I subtypes are essential for controlling lymph-borne viral infections after dissemination, with IFN-β playing a critical role in survival. **A-K**) The indicated mice were infected in the footpad with specified doses of ECTV (**A, E-H)** or ECTV Δ166 **(K**) for survival or 3,000 pfu of ECTV for quantifying titers in organs by plaque assay (**B-D, I-J**). Data are from two (**A-E, I-K**) or five (**F**) independent experiments combined. **G** and **H** correspond to one representative experiment. P values are compared with wild-type mice or the indicated groups using the Log-rank (Mantel-Cox) test for survival or t-test with Welch’s correction for viral titers experiments, in which each symbol represents an individual mouse. The dotted lines indicate the plaque assay detection limit (n.d.: not detected). **L**) WNV intra-host dissemination after subcutaneous footpad inoculation ^44^ (created with BioRender). **M-N**) Survival of the indicated mice infected in the footpad with 1,000 pfu of WNV (**M**) or KUNV (**N**). Data combined from three (**M**) or four (**N**) independent experiments. P values are compared with wild-type mice or the indicated groups using the Log-rank (Mantel-Cox) test.

Like *Ifna^Δ3^*, *Ifna^Δ6^*, and *Ifna4^−/−^* mice (mice only deficient in C&II IFN-α4, **Fig. 1C**) were as resistant to ECTV as controls at doses of 3,000 and 30,000 ECTV pfu (**Fig. 2F-G).** Different from *Ifna^Δ3^* and *Ifna^Δ6^* mice, which are deficient only in II IFN-I subtypes, a majority of *Ifnb1^−/−^* and to a greater extent *Ifna4,b1^−/−^*mice, which are deficient in both C&II IFN-I subtypes, succumbed to 3,000 and to 300 pfu ECTV (**Fig. 2F, H)**. Thus, C&II IFN-β plays a critical role in protecting from ECTV lethality, and C&II IFN-α4 enhances this resistance but is not required if IFN-β is present. Of note, ECTV was 100% lethal to *Ifnar1^−/−^* males and females, but more lethal to *Ifnb1^−/−^* males than females (**Sup. Fig. 1A-B**). Thus, some IFN-α subtypes may provide additional protection to females but not to males, supporting the notion that the IFN-I response contributes to sex differences in resistance to viral infection ^38^.

At five dpi, *Ifnar1*^−/−^, *Ifna4,b1*^−/−^, *Ifnb1*^−/−^, *Ifna4*^−/−^, and *Ifna^Δ6^* mice had higher viral loads than wild-type mice in their spleens, but only *Ifnar1*^−/−^ and *Ifna4,b1*^−/−^ mice had higher virus loads in their livers (**Fig. 2I**). At seven dpi, *Ifnar1*^−/−^, *Ifna4,b1*^−/−^, *Ifnb1*^−/−^, and *Ifna4*^−/−,^ but not *Ifna^Δ6^* mice, had higher virus loads than wild-type mice in their spleens and livers (**Fig. 2J**). Thus, at relatively late stages of infection and consistent with the survival data, the two C&II IFN-I subtypes, IFN-β and IFN-α4, are individually more important in controlling viral loads in organs during the systemic phase of infection than six II IFN-αs combined.

The ECTV protein EVM166 is an IFN-I decoy receptor ^39,40^ that binds IFN-β and IFN-α, ^41^ but only blocks IFN-α activity ^33^. ECTV deficient in EVM166 (ECTV Δ166) is highly attenuated but regains full lethality in *Ifnar1^−/−^* mice ^33,42^. Different to *Ifnar1^−/−^* mice, *Ifna4,b1*^−/−^, *Ifnb1*^−/−^, and *Ifna^Δ6^* mice were fully resistant to ECTV-Δ166 (**Fig. 2K**). Thus, within a context where ECTV is unable to block IFN-α, II IFN-αs suffice to control ECTV without the need for C&II positive feedback.

Because ECTV encodes a specific IFN-α inhibitor, the predominant role of C&II IFN-β in protecting the liver and resisting ECTV lethality could be unique to wild-type ECTV. Thus, we next analyzed the role of IFN-I subtypes in resistance to West Nile Virus (WNV), another lymph-borne virus.

WNV is a positive-sense single-stranded RNA flavivirus transmitted by mosquitoes. It can cause encephalitis and death in humans ^43^. WNV is also infectious to many other species, including mice ^44^. As with ECTV, WNV natural infection can be mimicked in mice by footpad inoculation. Despite its demonstrated trafficking from the skin to the dLN ^45,46^, pathogenic WNV strains, such as New York 2000 (NY2000, herein WNV)^46^, spread to the bloodstream as early as 2 dpi, bypassing the gatekeeping role of the innate immune response in the dLN. After systemic dissemination, WNV replicates in the spleen, brain, and gut (**Fig. 2L**). IFNAR is required to restrict viral replication in the brain ^47,48^ and the gut ^49^, and to resist lethality ^50,51^. Previous studies have shown that both *Ifnb1*^−/−^and wild-type mice treated with anti-IFN-α blocking antibodies are susceptible to WNV lethality ^47,51^.

We infected various IFN-I subtype-deficient mice with 1,000 pfu of WNV (**Fig. 2M**) or the naturally occurring attenuated WNV Kunjin strain (herein KUNV) (**Fig. 2N**), which is lethal to *Ifnar1^−/−^*but not to wild-type mice ^52^. Consistent with the literature ^47,52^, 50-60% of wild-type mice survived WNV infection, and all survived KUNV infection. Control *Ifnar1*^−/−^ mice quickly perished from both viruses. *Ifna4*^−/−^, *Ifna^Δ3^*, and *Ifna^Δ6^* mice were as resistant to WNV and KUNV as wild-type mice. Like with ECTV, *Ifnb1*^−/−^, and to a greater extent *Ifna4,b1*^−/−,^ were highly susceptible to WNV lethality. Moreover, ∼29% of *Ifnb1*^−/−^ and ∼53% *Ifna4,b1*^−/−^mice succumbed to KUNV. Hence, C&II IFN-β deficiency has drastic effects, while partial II IFN-α does not break resistance to lymph-borne DNA or RNA viruses’ lethality. Moreover, C&II IFN-α4 is not essential by itself, but supplements IFN-β to enhance resistance.

### 3. The NF-κB binding site in the Ifnb1 promoter and enhanceosome assembly are not essential for IFN-β production but are important in resisting ECTV and WNV lethality

Because IRF7 regulates all II IFN-I subtypes, IFN-β’s exceptional role in protection could be associated with its unique transcriptional regulation by constitutive TFs. We have previously shown that *Nfkb1*^−/−^ mice, but not *Irf3*^−/−^mice, are highly susceptible to ECTV lethality ^30,34^. However, *Nfkb1*^−/−^ mice lack lymph nodes and are impaired in the induction of multiple innate immune pathways ^53–55^. ECTV-infected iMOs, the major producers of IFN-I in the dLN at 2-3 dpi ^30^, upregulate *Nfkb1*, but not *Irf7*, *Irf3,* or *Jun* ^31^, and *Ifnb1’s* transcription depends on intrinsic NF-kB expression ^34^. Thus, to prevent NF-κB binding to the IFN-β’s enhancer, we partially deleted PRDII (*Ifnb1*^ΔPRDII^ mice) ^27^ (**Fig. 3A**). This deletion should also prevent HMG I(Y) binding to this region of the enhancer and enhanceosome assembly ^9,10,13^.

**Figure 3.**
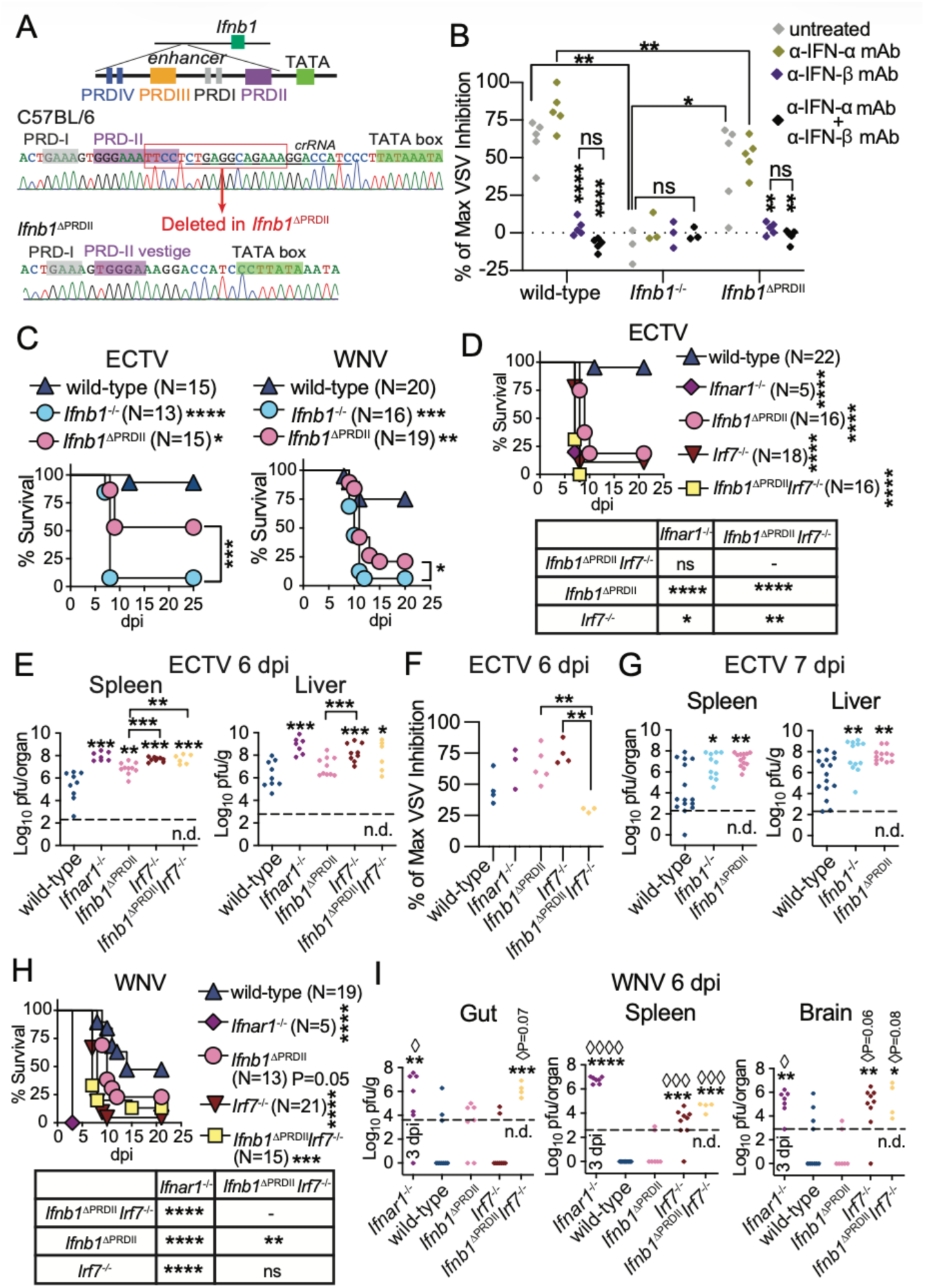
The NF-κB binding site in the Ifnb1 promoter and enhanceosome assembly are not essential for IFN-β production but are important in resisting ECTV and WNV lethality. **A**) Diagram of the *Ifnb1* enhancer in wild-type and in *Ifnb1*^ΔPRDII^ mice ^27^. **B**) Sera obtained from the indicated mice at seven dpi with ECTV were pre-treated or not with the indicated mAbs, and the IFN-I bioactivity was quantified by inhibition of VSV replication in mouse L929 cells. **C**) Survival of the indicated mice infected in the footpad with 3,000 pfu of ECTV or 1,000 pfu of WNV. The data correspond to three ECTV or two WNV independent experiments combined. **D**) Survival of the indicated mice infected in the footpad with 3,000 pfu of ECTV. **E**) ECTV titers in spleens and livers were determined by plaque assay at six dpi. **F**) The IFN-I bioactivity in sera from the indicated mice at six dpi with ECTV was quantified as in **B**. **G**) ECTV titers in spleens and livers were determined by plaque assay at seven dpi. **H**) Survival of the indicated mice infected in the footpad with 1,000 pfu of WNV. **I**) WNV titers in spleens, brains, and duodenum portions of the guts were determined by plaque assay at six dpi. The data correspond to one experiment (**B**), or two (**F**, **H**, **I**) or three (**D**, **E**, **G**) independent experiments combined. For **C**, **D,** and **H**, P values are compared with wild-type mice or the indicated groups using the Log-rank (Mantel-Cox) test. For **B**, **E-G,** and **I**, each diamond symbol depicts an individual mouse, and P values are compared with untreated controls (**B**), wild-type mice (**E**, **G**, **I**), or the indicated groups by t-test with Welch’s correction (asterisks) or by contingency Fisher test (◊ symbols). Dashed line indicates the plaque assay detection limit (n.d.: not detected).

To test whether *Ifnb1* could still be produced in *Ifnb1*^ΔPRDII^ mice despite disrupting NF-κB binding and enhanceosome assembly, we used a vesicular stomatitis virus (VSV) protection assay in the presence of blocking IFN-β or IFN-α monoclonal antibodies (mAbs) to determine IFN-β and IFN-α bioactivity in sera of mice at seven dpi with 3,000 pfu ECTV ^56^ (**Fig. 3B**). IFN-β was the predominant bioactive IFN-I in *Ifnb1*^ΔPRDII^ mice, albeit at reduced levels than in wild-type controls and was absent in *Ifnb1^−/−^* mice. Thus, *in vivo*, NF-kB binding to PRDII is important but not essential for systemic IFN-β production. Moreover, the data indicate that other TFs, such as IRF7, can upregulate *Ifnb1* in the absence of a complete enhanceosome. Consistent with the partial systemic reduction in IFN-β, *Ifnb1*^ΔPRDII^ mice were more susceptible to ECTV and WNV lethality than wild-type mice but less susceptible than *Ifnb1*^−/−^ mice (**Fig. 3C**). Therefore, C&II IFN-β regulation by constitutive NF-κB is more important to survive ECTV and WNV than up to six II IFN-α subtypes (compare **Fig. 3C** with **Fig. 2A, 2F, 2M**). Contrary to the current paradigm ^9,10,13,54^, IFN-β production can still proceed in the absence of a full enhanceosome.

IRF7 is an ISG required for the positive-feedback induction of all C&II and II IFN-I subtypes^17^, and is important in resisting ECTV and WNV lethality ^34,48^. Thus, we crossed *Ifnb1*^ΔPRDII^ with *Irf7*^−/−^ mice. The resulting *Ifnb1*^ΔPRDII^*Irf7*^−/−^ mice were as susceptible to ECTV lethality as *Ifnar1*^−/−^ mice. *Irf7*^−/−^ mice were slightly but more susceptible to ECTV lethality than *Ifnb1*^ΔPRDII^ mice (P=0.0122) (**Fig. 3D)**. At six dpi, the spleens of *Ifnb1*^ΔPRDII^, *Irf7*^−/−^ and *Ifnb1*^ΔPRDII^*Irf7*^−/−^ mice, and the livers of *Irf7*^−/−^ and *Ifnb1*^ΔPRDII^*Irf7*^−/−^ mice had higher viral loads than those of wild-type mice (**Fig. 3E**). At this time, and despite the increased virus loads, the systemic bioactive IFN-I levels were lower in *Ifnb1*^ΔPRDII^*Irf7*^−/−^ than in *Ifnb1*^ΔPRDII^ or *Irf7*^−/−^ mice, and trended lower than in wild-type mice, while those in *Ifnar1^−/−^*, *Ifnb1*^ΔPRDII^ and *Irf7*^−/−^ mice trended higher than in control mice, likely driven by the increased virus loads (**Fig. 3F**). Given that all or most of the bioactive IFN-I in the sera of ECTV-infected mice is IFN-β (**Fig. 3B**), these data demonstrate that after ECTV infection, NF-κB and IRF7 can independently induce IFN-β production without the need for the assembly of a complete enhanceosome, and that NF-κB and IRF7 are responsible for most of the IFN-β production during the systemic phase of ECTV infection. Of note, at seven dpi, *Ifnb1*^ΔPRDII^ and *Ifnb1*^−/−^ mice had higher viral loads in their spleens and livers than wild-type mice (**Fig. 3G**). Hence, controlling ECTV loads in organs requires constitutive NF-κB induction of IFN-β at a relatively late stage of systemic infection, when disease signs typically appear.

Compared to wild-type, *Irf7*^−/−^ mice were more susceptible than *Ifnb1*^ΔPRDII^ (P<0.001) mice, as susceptible as *Ifnb1*^ΔPRDII^*Irf7*^−/−^ mice, and less susceptible than *Ifnar1*^−/−^ mice to WNV lethality (**Fig. 3H**). At six dpi, most *Irf7*^−/−^ and *Ifnb1*^ΔPRDII^*Irf7*^−/−^ mice had higher viral loads in the spleen and brain, while *Ifnb1*^ΔPRDII^*Irf7*^−/−^ but not *Irf7^−/−^* mice had high viral loads in the gut, suggesting that NF-κB induced-IFN-β can independently control WNV in the gut. In contrast, viral loads in *Ifnb1*^ΔPRDII^ mice were similar to those in wild-type mice, with most mice having undetectable viral loads in the spleen and brain (**Fig. 3I**).

Collectively, **Fig. 3** indicates that IRF7 and NF-kB synergize to protect from lymph-borne virus lethality, that IFN-I production *via* constitutive NF-κB or inducible IRF7 vary in their ability to control virus loads in different organs, and that IFN-β production can occur without the assembly of a full enhanceosome.

### 4. IFN-α curbs ECTV systemic spread, while IFN-β restricts viral replication in the liver and prevents lethality

Next, we created mice deleted of all IFN-α genes (*Ifna*^−/−^) mice (**Fig. 4A**). *Ifna*^−/−^ mice were slightly but more susceptible than wild-type mice to ECTV lethality, but much less than *Ifnb1*^−/−^ mice (**Fig. 4B**).

**Figure 4.**
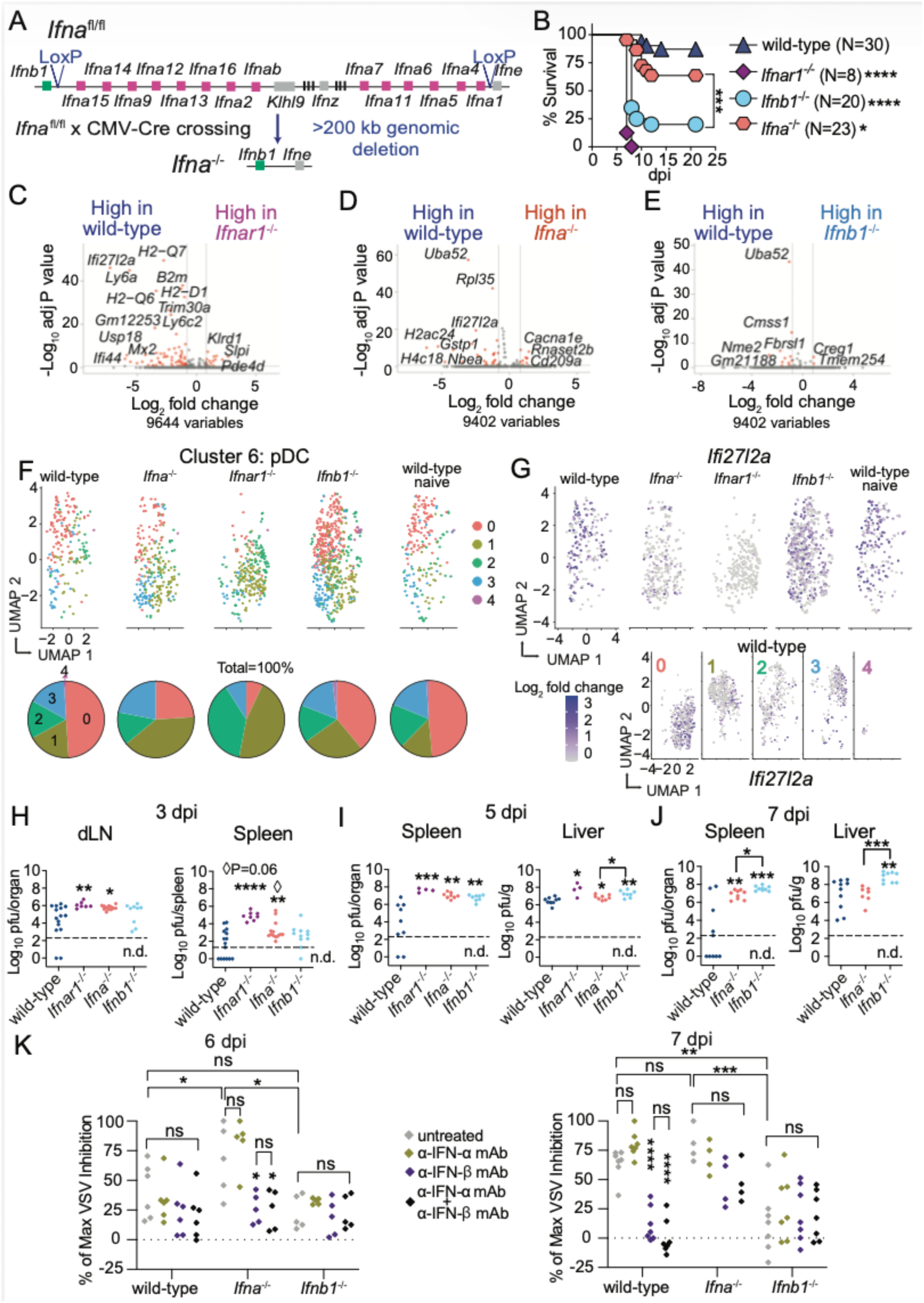
IFN-α curbs ECTV systemic spread, while IFN-β restricts viral replication in the liver and prevents ECTV lethality. **A**) Diagram depicting the germline deletion of all *Ifna* genes in *Ifna*^−/−^ mice produced by crossing *Ifna*^fl/fl^ mice ^27^ with CMV-Cre mice. **B**) Survival of mice infected in the footpad with 3,000 pfu of ECTV. The data correspond to three independent experiments combined. **C-E**) Volcano plots showing differentially expressed genes of clusters annotated as pDCs (SiglecH^+^Ly6C^+^CD4^+^CD11c^+^CD11b^−^Ly6G^−^NK1.1^−^CD103^−^) comparing indicated groups (red dots: Log2 fold change=0.8 and adj P<0.2, gray dots: unchanged). **F**) UMAP showing re-clustering of pDCs and pie charts of percentages of individual sub-clusters in each sample. **G**) UMAP showing *Ifi27l2a* gene expression in pDCs from each sample (top) and in pDCs sub-clusters from infected wild-type mice (bottom). **H-J**) ECTV titers in popliteal draining lymph nodes, spleens, and livers determined by plaque assay at three (**H**), five (**I**), and seven (**J**) dpi. Data correspond to three (**H**) or two (**I, J**) independent experiments combined. **K**) Sera obtained from the indicated mice at six (left) or seven (right) dpi with ECTV were pre-treated or not with the indicated mAbs, and the IFN-I bioactivity was quantified by inhibition of VSV replication in mouse L929 cells. The data correspond to two independent experiments combined. For **B**, the P values are compared with wild-type mice or the indicated groups using the Log-rank (Mantel-Cox) test. For **H-J**, each symbol depicts an individual mouse. The P values are compared with wild-type mice (**H-J**), to untreated controls (**K**), or to the indicated groups by t-test with Welch’s correction. Dashed line indicates the plaque assay detection limit (n.d.: not detected).

It is widely believed that basal levels of IFNAR signaling poise steady-state myeloid cells to respond rapidly to infections ^31,57^. To test whether IFN-α- or IFN-β-deficiency affects the very early innate immune response to ECTV as it reaches the dLN ^58^, we FACS-sorted live innate immune cells from the dLNs of wild-type uninfected, or wild-type, *Ifnar1*^−/−^, *Ifna*^−/−^, and *Ifnb1*^−/−^ mice at one dpi with ECTV. We performed cellular indexing of transcriptome and epitope single cell sequencing (CITEseq) (**Sup. Fig. 2A**). Using Uniform Manifold Approximation and Projection (UMAP) clustering of cells, we identified nineteen distinct cell clusters across all the samples (**Sup. Fig. 2B**). Based on protein and transcript expression, each cluster was assigned to a specific cell population (**Table S1**). The absence of IFNAR, IFN-α, or IFN-β did not affect the innate immune cell composition of the dLN at one dpi (**Sup. Fig. 2C, Table S1**). Using plaque assays, confocal microscopy, and qPCR after cell sorting, we previously showed that at one dpi, ECTV is already present in the dLN within a few migratory dendritic cells (mDCs) ^32,59,60^. However, with CITE-Seq, we did not detect ECTV transcripts in any cell population, probably reflecting that only a few cells are infected with ECTV at this early time point. When comparing wild-type cells from naïve and infected mice, we observed few infection-induced genes in these cell populations (**Table S2**), suggesting that at one dpi, ECTV does not yet induce a robust innate immune response in the dLN.

However, we observed transcriptional changes in plasmacytoid DCs (pDCs) (**Fig. 4C-E**, **Table S3**), and to a lower extent in NK cells, mDCs, conventional DCs (cDC), and iMOs (**Sup Fig. 2E-G**), from IFN-I-defective compared to IFN-I-sufficient infected mice. Compared to wild-type pDCs, *Ifnar1*^−/−^ *Ifna*^−/−^ and *Ifnb1*^−/−^ pDCs had respectively lower expression of 65, 49, and 11 gene transcripts (Log_2_ fold change > 0.8, adjusted P < 0.2). *Irf7* was downregulated in *Ifnar1*^−/−^ pDC but not in *Ifna*^−/−^ or *Ifnb1*^−/−^ pDCs, indicating that these cytokines redundantly prime pDCs for IFN-I transcription. *Irf9* was downregulated in *Ifnar1*^−/−^ (adjusted P = 0.004) and *Ifna*^−/−^ (P = 0.0001) pDCs but not in *Ifnb1*^−/−^ pDCs, indicating that IFN-α may specifically prime pDCs for ISG upregulation. However, these observations require further validation, especially for *Irf9* downregulation in *Ifna*^−/−^ pDCs, as it did not meet the false discovery test threshold of 0.2 adjusted P-value, which was used as the baseline for our studies. We identified two genes that were downregulated in *Ifnar1*^−/−^, *Ifna*^−/−^, and *Ifnb1*^−/−^ pDCs compared to wild-type pDCs: *Ifi27l2a* and *Nbea*. IFI27L2a is an ISG important for controlling WNV infection ^61^. NBEA has been associated with neurological diseases, and no role for anti-viral responses has been described so far. By re-clustering pDCs to identify IFN-I responder cells, we observed five pDC sub-clusters similarly distributed between pDCs from naïve and infected wild-type mice (**Fig. 4F**). Clusters 0 and 3 were greatly depleted in pDCs from *Ifnar1*^−/−^ mice, and cluster 0 was depleted in pDCs from *Ifna*^−/−^ mice. In addition, *Ifi27l2a* transcription was predominantly localized in clusters 0 and 3 and greatly reduced in *Ifnar1*^−/−^ and *Ifna*^−/−^ pDCs (**Fig. 4G**). Overall, IFN-α, and to a much lower degree IFN-β, prime pDCs in the dLN as early as one dpi. Even though pDCs are not required to survive ECTV infection ^30^, they may contribute to curbing ECTV replication in the dLN.

At three dpi, *Ifna*^−/−^ but not *Ifnb1*^−/−^ mice presented higher viral loads in the dLN compared to controls (**Fig. 4H**, **Fig. 1F**). Like the other IFN-I knockout strains (**Fig. 1F**), *Ifna*^−/−^ mice presented faster dissemination to the spleen at three dpi compared to wild-type mice. *Ifnb1^−/−^* and *Ifna*^−/−^ mice had higher ECTV loads than wild-type mice in the liver and spleen at five dpi, although the virus loads in the liver were slightly lower in *Ifna^−/−^* than in *Ifnb1^−/−^* mice (**Fig. 4I**). At seven dpi, ECTV titers in the spleens of *Ifnb1^−/−^* and *Ifna*^−/−^ mice were still higher than in controls, but only *Ifnb1^−/−^* mice had higher ECTV titers in the liver (**Fig. 4J**).

At six dpi, few wild-type mice presented bioactive IFN-I in their sera (**Fig. 4K, left)**, which peaked at seven dpi in this strain (**Fig. 4K, right**). Notably, the levels of bioactive IFN-I were higher at six dpi and similar at seven dpi in *Ifna^−/−^* mice compared to wild-type mice. At six and seven dpi, the levels of bioactive IFN-I in *Ifnb1^−/−^*mice were at basal levels and lower than in wild-type and *Ifna^−/−^* mice. Also, blockade with anti-IFN-β and anti-IFN-β+IFN-α but not with anti-IFN-α mAbs reduced the IFN-I bioactivity in sera from wild-type at seven dpi and from *Ifna^−/−^*mice at six dpi. At seven dpi, anti-IFN-β and anti-IFN-β+IFN-α mAb treatment did not fully block the IFN-I bioactivity in sera from *Ifna^−/−^* mice. Given the high levels of antibody used in the assay (20 μg/well), these data suggest that at seven dpi with ECTV, *Ifna*^−/−^ mice sera contain compensatory levels of IFN-β and possibly other cytokine(s) that suppress VSV replication, such as IFN-γ ^62^. This indicates an increase in IFN-β production to compensate for the absence of IFN-α in *Ifna^−/−^* mice but no compensatory IFN-α activity in *Ifnb1^−/−^*mice.

Interestingly, at six and seven dpi with ECTV, the sera from wild-type female mice contained more bioactive IFN-I than the sera from wild-type male mice (**Sup Fig. 3A-B**). At six dpi with ECTV, the sera from *Ifna*^−/−^ female mice had higher IFN-I activity than the sera from male mice. However, at seven dpi, the sera from *Ifna*^−/−^ mice of both sexes had similarly high levels of IFN-I. Male and female *Ifnb1*^−/−^ mice had low levels of IFN-I bioactivity at six and seven dpi, but slightly higher in females than in males at seven dpi. Therefore, increased mousepox susceptibility and higher viral burden in males compared to females (**Sup Fig. 1, Fig. 2C**) correlated with lower levels of systemic IFN-I.

Overall, **Fig.4** shows that IFN-α, and to a lesser extent IFN-β, primes pDCs and other innate immune cells in the dLN, curbs ECTV spread, and controls replication in organs at the initial stages of the systemic infection. However, at later stages, IFN-β but not IFN-α is required to restrict ECTV replication in the liver, the main target organ for ECTV lethality.

### 5. IFN-α and IFN-β non-redundantly protect from WNV infection by restricting viral replication in the brain at different times post-infection

We showed above that IFN-β is more important than multiple IFN-α subtypes in resisting WNV lethality. Yet, antibody-blocking experiments in mice indicated that resistance to WNV lethality requires both IFN-α and IFN-β ^56^. To definitively determine and compare the contribution of IFN-β and IFN-α in resistance to WNV, we used *Ifna*^−/−^ mice. *Ifna*^−/−^ mice were 100% susceptible to WNV lethality, which was higher than for *Ifnb1*^−/−^ mice. While *Ifna*^−/−^mice succumbed later than *Ifnar1*^−/−^ mice (P<0.0001, mean time of death at 8 dpi), they died earlier than *Ifnb1^−/−^* mice (mean time of death at 11 dpi) (**Fig. 5A**). These data indicate that IFN-α plays a more significant role and contributes earlier to surviving WNV infection than IFN-β. This is contrary to the expectation of IFN-β as “early” and most of the IFN-α subtypes as “late” IFN-I subtypes. Of note, *Ifna*^−/−^ mice are deficient in IFN-ζ and KLHL9. However, the high susceptibility of *Ifna^−/−^* mice cannot be attributed to these deficiencies because *Ifna*^β6^ mice also lack IFN-ζ and KLHL9 but are as resistant to WNV lethality as wild-type mice (**Fig. 2M**).

**Figure 5.**
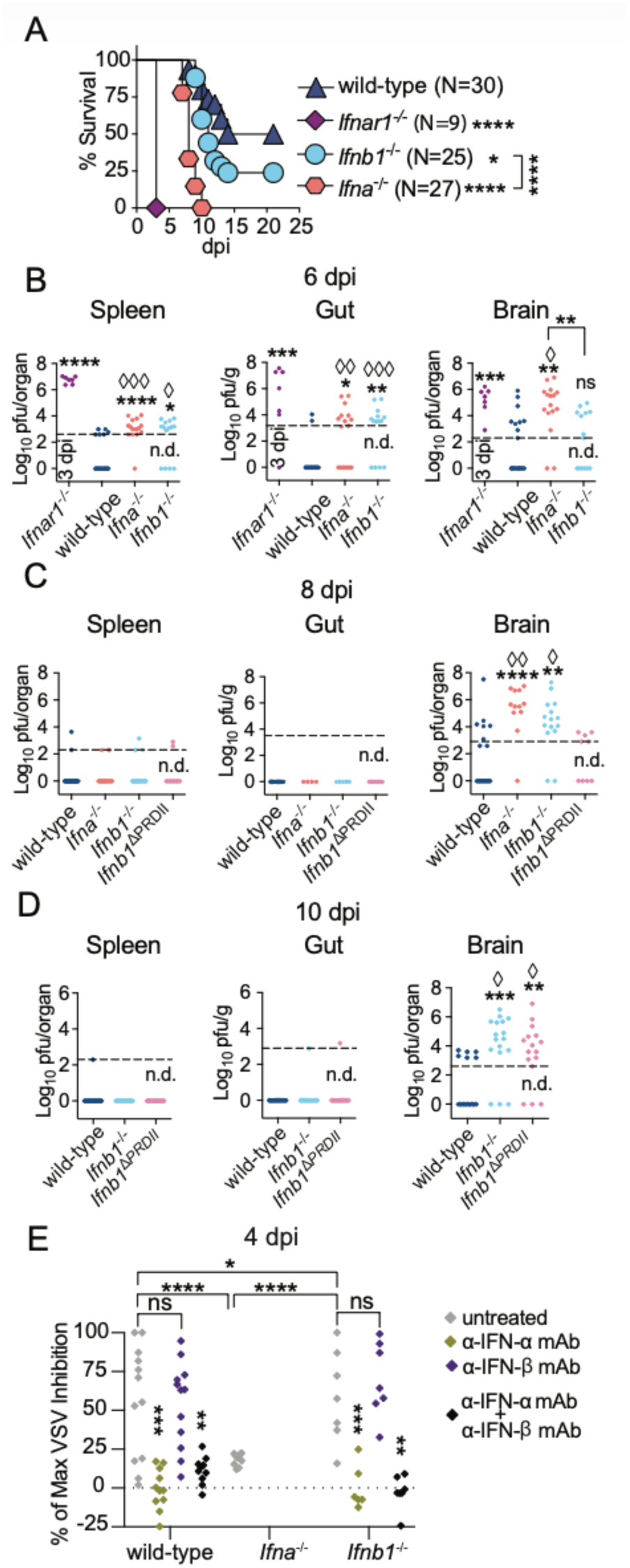
IFN-α and IFN-β non-redundantly protect from WNV infection by restricting viral replication in the brain. **A)** Survival of mice infected in the footpad with 1,000 pfu of WNV. The data correspond to three independent experiments combined. **B-D)** WNV titers in spleens, brains, and duodenum portions of the gut were determined by plaque assay at six (**B**), eight (**C**), and ten (**D**) dpi. The data correspond to three independent experiments combined. **E)** Sera obtained from mice at four dpi with WNV were pre-treated or not with the indicated mAbs, and the IFN-I bioactivity was quantified by inhibition of VSV replication in mouse L929 cells. The data correspond to two independent experiments combined. In **A**, the P values are compared with wild-type mice or the indicated groups using the Log-rank (Mantel-Cox) test. In **B-E**, each symbol depicts an individual mouse. The P values are compared with wild-type mice (**B-D**) or untreated controls (**E**) or the indicated groups using the t-test with Welch’s correction (asterisks) or using the contingency Fisher test (◊ symbols).

At six dpi, *Ifna*^−/−^ and *Ifnb1*^−/−^ groups had higher frequencies of mice with detectable WNV and overall higher WNV loads in the spleen (respectively P <0.0001 and <0.0001) and guts (respectively P = 0.0132 and 0.0233) than wild-type mice but lower than in moribund *Ifnar1*^−/−^ mice at three dpi (**Fig. 5B**). This suggests that restriction of WNV systemic dissemination to spleen and gut requires the combined action of IFN-α and IFN-β. Also at six dpi, but in the brain, *Ifna*^−/−^ mice, but not *Ifnb1*^−/−^ mice, presented higher viral loads and a higher frequency of animals with detectable WNV compared to controls, and not significantly different from WNV loads in *Ifnar1*^−/−^ mice at three dpi. This indicates that IFN-α plays an earlier role than IFN-β in protecting the brain from WNV replication, further contradicting the view that IFN-β is an early subtype and most IFN-αs are late.

At eight (**Fig. 5C**) and ten (**Fig. 5D**) dpi, WNV was no longer detected in the gut and only few mice sill had detectable virus in the spleen in wild-type, *Ifna*^−/−,^ *Ifnb1*^−/−,^ and *Ifnb1*^ΔPRDII^ mice, suggesting that in the gut and spleen WNV can be controlled by IFN-β, IFN-α, or independently of IFN-I. In contrast, *Ifna*^−/−^ and *Ifnb1*^−/−^ mice had higher WNV burdens and a higher frequency of mice with WNV in the brain than wild-type and *Ifnb1*^ΔPRDII^ mice (**Fig. 5C**).

At 10 dpi, when all *Ifna^−/−^* mice were already dead, *Ifnb1*^−/−^ and *Ifnb1*^ΔPRDII^ mice had higher viral loads than wild-type mice in their brains (**Fig. 5D**). Notably, the timing of increased viral loads in the brains of *Ifna*^−/−^ (six dpi), *Ifnb1*^−/−^ (eight dpi), and *Ifnb1*^ΔPRDII^ (ten dpi) mice preceded their mean time of death of eight dpi for *Ifna*^−/−^, ten dpi for *Ifnb1*^−/−^ and eleven dpi for *Ifnb1*^ΔPRDII^ mice. These data suggest that first IFN-α, then NF-κB-independent IFN-β, and subsequently NF-κB-dependent IFN-β protect the brain from WNV replication.

The IFN-I bioactivity in the sera of WNV-infected mice peaks at ∼4 dpi ^56^. At this time, the sera from *Ifnb1*^−/−^ mice had higher levels of IFN-I bioactivity than sera from wild-type mice, whereas those in the sera from *Ifna^−/−^*mice were basal (**Fig. 5E**). Moreover, the IFN-I bioactivity in sera from wild-type and *Ifnb1^−/−^* mice was due to IFN-α because it was blocked with anti-IFN-α or anti-IFN-α+anti-IFN-β, but not with anti-IFN-β mAbs.

Overall, **Fig. 5** indicates that, unlike ECTV infection, most IFN-I bioactivity in the sera from WNV-infected mice is due to IFN-α, rather than IFN-β. Moreover, *Ifna*^−/−^ mice do not produce compensatory systemic IFN-β, while *Ifnb1^−/−^* mice compensate with increased IFN-α bioactivity. Our data also indicate that one or a few IFN-α subtypes, not deleted in *Ifna*^Δ3^ or *Ifna*^Δ6^ mice, or a specific threshold of IFN-α is required to restrict viral loads in the brain before six dpi and reduce WNV lethality. Moreover, although IFN-β and PRDII provide a survival advantage, mice do not have systemic IFN-β at four dpi with WNV. This suggests that systemic IFN-β distribution occurs late after WNV infection or, more likely, that for WNV, IFN-β operates in tissues rather than systemically, and is time-restricted to specific organs, such as the brain, after six dpi.

## Discussion

It is well established that IFN-I binding to IFNAR initiates a signaling cascade that is essential for curbing the disease and lethality caused by multiple pathogens ^63–66^. However, the overall and temporal roles of the multiple IFN-I subtypes, and their regulation by constitutive and IFN-I-inducible TFs in anti-viral defense *in vivo*, remain unclear. A key finding from our studies is that IFN-α and IFN-β play distinct roles in the peripheral and systemic phases of viral infection. Our finding that all tested IFN-I-deficient strains exhibit defective ECTV containment in the lymph nodes suggests that the high multiplicity of IFN-I subtypes maximizes positive feedback signaling, thereby restricting systemic viral dissemination. We also demonstrate that multiple IFN-α subtypes are dispensable for survival during ECTV and WNV infections. Notably, a genetic study of healthy individuals worldwide revealed a 34% homozygosity frequency for a null allele of IFN-α10, and that some IFN-α subtypes have high frequencies of nonsynonymous variants ^67^. This suggests that, as we observed in *Ifna^Δ3^* and *Ifna^Δ6^* mice infected with ECTV and WNV, some IFN-α subtypes may be dispensable for overall antiviral resistance in humans. Together, our results in mice with partial deficiency in IFN-α subtypes suggest that the physiological advantage of increasing the number of IFN-I genes is to contain systemic viral spread, rather than to enhance survival to acute viral infections. We also showed that IFN-α is more critical than IFN-β for containing ECTV and activating pDC in the dLN as early as one dpi. Unlike most cells, pDCs constitutively express IRF7 ^68^. Thus, our results indicate cell- and tissue-specific constraints on IFN-I production, and that peripheral cells may depend on IRF7-IFNα interactions to prevent viral diseases.

Our finding that, individually, IFN-β is by far the most effective IFN-I subtype for resisting two dissimilar lymph-borne viruses correlates with IFN-β’s unique promoter and higher affinity for IFNAR ^19,20^, which are conserved between humans and mice. We also demonstrated that IFN-α is essential for survival during WNV infection, but not during ECTV infection. Yet, deletion of six IFN-α subtypes does not increase the lethality of WNV, and at least IFN-α1, -α2, -α4, -α5, -α6, -α7, -α11, -α16, and -αab are not individually required for survival to WNV infection. It remains unresolved whether survival to WNV infection requires a minimum number of IFN-α subtypes, regardless of subtype identity, or whether one or more specific subtypes, present in *Ifna^Δ3^* and *Ifna^Δ6^*, are particularly critical.

Another striking finding was that sera from ECTV-infected mice contain bioactive IFN-β but not -α, whereas, as others have shown^51^, sera from WNV-infected mice contain bioactive IFN-α but not -β. It is unclear whether the sera of ECTV-infected mice lack IFN-α or are fully blocked by circulating EVM166^33^. In the case of WNV, which is not known to block any IFN-I subtype directly, it is possible that WNV selectively induces systemic production of IFN-α, or that it selectively inhibits systemic *Ifnb1* transcription, such as by inducing sumoylation of the *Ifnb1* distal enhancer ^69^. The subtype divergence in IFN-I bioactivity during ECTV and WNV infections may also reflect differences in the signaling pathways whereby they induce IFN-I. While IFN-I expression and maximal survival to ECTV and WNV require cGAS and IRF7 ^35,70–72^, WNV also requires RIG-I and IRF3 ^34,52,73,74^. Yet, given that IFN-β is necessary to restrict viral replication in the brain and to support survival at later stages of infection, WNV must induce IFN-β production, at least locally in the brain, at later stages of infection.

Previous studies have clearly established that deletion of nucleotides absent in *Ifnb1*^βPRDII^ mice prevent the binding of p65 and HMG I(Y) to PRDII and enhanceosome assembly ^75,76^. Thus, our results with *Ifnb1*^βPRD^ ^II^ mice show that PRDII is important for optimal IFN-β production and survival against ECTV and WNV infection. However, *Ifnb1*^βPRDII^ mice have IFN-β in the serum following ECTV infection and are less susceptible to ECTV or WNV lethality than *Ifnb1^−/−^* mice. These data indicate that NF-kB plays an important role in optimal IFN-β production and survival against ECTV and WNV, but that IFN-β production can proceed and protect mice from lethality without the assembly of a full enhanceosome. Notably, mice deficient in PRDII and IRF7 are as susceptible to ECTV as *Ifnar1^−/−^* mice, suggesting that these two TFs are the essential mediators of protective IFN-I production in response to ECTV. We previously showed that IRF3 is dispensable^30,34^; whether AP-1 is required remains unresolved.

Our data also show that, despite increased systemic ECTV titers in *Ifna4^−/−^*, *Ifna^Δ6^*, and *Ifna^−/−^* mice, IFN-β is sufficient to control excessive ECTV replication in the liver and to afford survival. Given that the peak of bioactive IFN-β in the sera coincides with the key seven dpi time-point when viral loads increase in the liver of *Ifnb1*^−/−^ mice, the bioactive IFN-β in the liver could be produced by hematopoietic or parenchymal liver cells.

Contrary to the current paradigm that constitutively-induced IFN-β is early and II-IFN-α subtypes are late, we found that WNV titers become high in the brains of *Ifna^−/−^*, *Ifnb1*^−/−,^ and *Ifnb1*^βPRDII^ mice at six, eight, and ten dpi, respectively, preceding the mean time of death of each strain. This suggests that IFN-I subtypes control virus titers in the brain and protect from death in different waves. First, by IFN-αs likely produced systemically, next by IFN-β produced by an NF-κB-independent mechanism likely within the brain, and then by IFN-β through an NF-κB-dependent mechanism likely also within the brain. The temporal need for NF-κB-independent and -dependent IFN-β production suggest a requirement for different cellular sources of IFN-β at different times.

Finally, while our data addresses critical issues regarding IFN-I subtypes and their control during acute lymph-borne viral infections, IFN-I also plays important roles in other infectious diseases, as well as in development, cancer, autoimmunity, and neurodegeneration ^77–80^. Thus, our findings may have broader implications, and our mouse models may help address other critical questions.

## Material and Methods

### Ethics Statement

All the procedures involving mice were carried out in strict accordance with the recommendations in the Eighth Edition of the Guide for the Care and Use of Laboratory Animals of the National Research Council of the National Academies. All experiments were approved by Thomas Jefferson University’s Institutional Animal Care and Use Committee under protocol 01727 “Innate Control of Viral Infections.”

### Reagents

All reagents used are listed in Supplemental Material, Box 1.

### Mice

Male and female mice used in experiments were 6 to 22 weeks old. C57BL/6NCrl mice were purchased from Charles River or bred at Thomas Jefferson University Laboratory Animal Facility. *Ifnar1*^−/−^ mice backcrossed to C57BL/6 as gifts from Dr. Thomas Moran (Mount Sinai School of Medicine, New York, NY). B6;129P2-Irf7tm1Ttg/TtgRbrc (*Irf7*^−/−^) mice were obtained from Dr. Tadatsugu Taniguchi ^68^. B6.C-Tg(CMV-cre)1Cgn/J mice were purchased from the Jackson Laboratory. *Ifnb1*^−/−^, *Ifna4,b1*^−/−^, *Ifna4*^−/−^, *Ifna*^β3^, *Ifna*^β6^, *Ifnb1*^ΔPRDII^ and *Ifna*^fl/fl^ mice were generated in our laboratory using CRISPR/Cas and iGONAD, as previously published ^27^. Briefly, CRISPR RNA and Cas9 protein were injected into the oviducts of superovulated, 0.7-day pregnant C57BL/6NCrl females and subsequently electroporated. Founders were screened by Sanger sequencing for on- and off-target genomic sites with a minimum cutting frequency threshold of 0.2, as determined by the CRISPOR tool ^81^. Selected founders were bred with C57BL/6NCrl mice. The resulting heterozygous offspring were used to generate homozygous genetically modified strains. *Ifnb1*^ΔPRDII^*Irf7*^−/−^ mice were made by crossing *Ifnb1*^ΔPRDII^ with *Irf7*^−/−^ mice. *Ifna*^−/−^ mice were made by crossing *Ifna*^fl/fl^ ^27^ with B6.C-Tg(CMV-cre)1Cgn/J (CMV-Cre) mice ^82^, which drives germline expression of Cre recombinase by the cytomegalovirus promoter. Colonies were bred at Thomas Jefferson University under specific pathogen-free conditions. Animals were housed in ventilated racks with a light/dark cycle and fed *ad libitum* on 5010 LabDiet (Quakertown, PA, USA).

### Viruses and infection

ECTV Moscow strain (ATCC VR-1374), green fluorescent protein (GFP)-expressing ECTV ^83^, and ECTV Δ166 ^33^ were propagated in tissue culture. Briefly, BS-C-1 cells (ATCC CCL-26) grown in supplemented DMEM media to 80-90% confluency were infected at a multiplicity of infection of 0.01, and viruses were harvested 4 to 5 days later before cell detachment. Stocks were titrated by plaque assay in BS-C-1 cells, as described below. KUNV (CH16532) was a generous gift from R. Tesh (World Reference Center of Emerging Viruses and Arboviruses, Galveston, TX), and WNV NY2000 (herein WNV) was a generous gift from Michael Diamond. KUNV stocks were propagated in BHK-21 cells (ATTC CCL-10). WNV and KUNV stocks were propagated in C6/36 Aedes albopictus cells (ATCC CRL-1660) and titrated in BHK-21 cells by determining the 50% tissue culture infective dose, as previously described ^84^. Vesicular stomatitis virus (VSV) Indiana strain (provided by J.T. Guo, Drexel Institute for Biotechnology and Virology Research, Doylestown, PA) was expanded in A9 cells (ATCC CRL-1811), and the viral yield was determined by a plaque assay on Vero cells (ATCC CCL-81), as previously described^33^.

### Infections

Mice were infected subcutaneously in the rear footpad with the indicated doses of plaque-forming units (PFU) of ECTV, ECTV-GFP, ECTV Δ166, WNV or KUNV. In survival studies, mice were monitored daily and sacrificed when illness resulted in inactivity, unresponsiveness to touch, limb paralysis, or more than 25% weight loss. Euthanasia was performed according to the 2013 edition of the AVMA Guideline for the Euthanasia of Animals.

### Virus titration

Entire or portions of organs were homogenized in 10% fetal bovine serum-supplemented DMEM media using a TissueLyzer II (Qiagen), and titers were determined by plaque assay on BS-C-1 cells for ECTV ^32^ or on BHK-21 for WNV ^85^. Shortly, BS-C-1 cells at 80-90% confluency or BHK21 cells at 50-60% confluency were infected with 10-fold dilutions of each virus sample for 2 hours in supplemented DMEM media. Infected cells were grown in 1% carboxymethyl cellulose overlay containing 2% fetal bovine serum-supplemented DMEM media. Virus plaques were quantified at five dpi for ECTV or at three to four dpi for WNV after fixation with 4% formaldehyde for 10 minutes and staining with 0.2% crystal violet dissolved in 20% methanol.

### Quantification of IFN-I transcription

As previously described ^31,32^, mice infected with ECTV-GFP in both rear footpads were euthanized at three dpi. Spleens from naïve mice were used for sorting naïve iMOs. Lymph nodes were treated with Liberase TM for 30 min, and single-cell suspensions of pooled lymph nodes were prepared using a 70 μm strainer. Cells were treated with anti-mouse CD16/CD32 (BioLegend - 0.25 μL per million of cells) for 15 min at 4°C and subsequently stained with surface antibodies for 20 min at 4°C. After washing, iMOs were sorted with a FACSAria™ II sorter at Thomas Jefferson University’s Flow Cytometry and Human Monitoring Shared Facility. Cells were sorted into TRIzol, and the RNA was purified using the RNA Clean and Concentrator (Zymo Research). RNA was used as template for cDNA synthesis with the High-Capacity cDNA Reverse Transcription Kit, and Quantitative PCR (qPCR) was performed, as previously described ^30,34^, using SYBR Green PCR Master Mix in a Thermocycler CFX96 Real-Time System (BioRad). Gene expression was normalized by *Gapdh* levels, and fold change induction was calculated relative to iMOs sorted from naïve mice.

### Quantification of IFN-I bioactivity

L929 cells (ATCC CCL-1) were treated with sera from infected animals overnight and infected with VSV for 24 hours. Cells were fixed with 4% formaldehyde for 10 minutes and stained with 0.2% crystal violet dissolved in 20% methanol. Cell viability was quantified by OD595 measurement and % VSV inhibition calculated as follow: (OD595_serum_ – OD595_no serum_) * 100/(OD595_mock_ – OD595_no serum_). The % maximum VSV inhibition was calculated to normalize the data, ensuring that the highest measurement was 100%. Sera was untreated or pre-incubated at 37°C for 1 hour with 20 μg of anti-mouse IFN-β (Leinco HDβ-4A7) and/or anti-mouse IFN-α (Leinco TIF-3C5) mAbs^51^. Recombinant mouse IFN-β, IFN-α1 or IFN-α3 (PBL assay science) were used as controls.

### CITEseq

For sorting live innate immune cells from popliteal dLNs, naïve mice or mice previously infected with ECTV in both rear footpads were euthanized at three dpi. Lymph nodes were processed as described for “*Quantification of IFN-I transcription”*, using low-binding nuclease-free and wide-bore pipette tips. After live/dead stain with Zombie Violet in serum-free conditions for 15 min at room temperature, cells were treated with anti-mouse CD16/CD32 (BioLegend - 0.25 μL per million of cells) for 15 min at 4°C. TotalSeq-B and fluorescent-labeled surface antibodies (Box 1) were added and incubated for 30 min at 4°C. Labeled cells were resuspended in 2% fetal bovine serum-supplemented phosphate buffered solution and sorted into 20% fetal bovine serum-supplemented RPMI media using EDTA-free sheath fluid on a BD Symphony S6 Sorter at Penn Cytomics and Cell Sorting Resource Laboratory, University of Pennsylvania. Cells were centrifuged, resuspended in RPMI media supplemented with 10% fetal bovine serum, and counted with trypan-blue exclusion (viability > 90%). Cells were loaded into a 10X Genomics Chromium chip for targeted cell recovery of 10,000 cells, then loaded into a Chromium X controller. Resultant single-cell barcoded gel beads-in-emulsion were processed according to 10X Genomics’ protocols for the construction of mRNA 3’Gene Expression and of antibody Cell Surface/Feature libraries. Quality control-approved libraries were sequenced on a Novaseq6000 Illumina sequencer by the Cancer Genomics Core at Thomas Jefferson University.

### Quantification, Statistical analysis, and sequencing data analysis

Data were analyzed with Prism 10 Software. Log-rank (Mantel-Cox) analysis was used for survival experiments. Unpaired Student’s t-test with Welch’s correction (asterisks) and contingency Fisher test (◊ symbols) were used as applicable for other experiments. In all figures, * or ◊ P < 0.05, ** or ◊◊ P < 0.01, *** or ◊◊◊ P < 0.001, **** or ◊◊◊◊ P < 0.0001. CITE-seq raw reads were processed and aligned with the 10X Genomics cellranger tool (version 8.0.1). To generate the working reference genome, we used RStudio (4.5.5.1) to combine the mouse GRCm39 mm10 (C57BL/6) reference genome with the GCA_000841905.1 ectromelia virus Moscow reference genome. Data from all subjects were combined in a single object. Annotated data were processed using the Seurat package (version 5.2.0) ^86^, running in RStudio (4.5.5.1), as previously described ^87–90^. Briefly, quality control and cell selection were performed by removing cells with more than 5% mitochondrial genes and retaining those with fewer than 6000 nCountRNA and between 200 and 70000 detected genes (nFeature_RNA). The data were normalized using the Seurat function “LogNormalize”. The highly variable features were detected using Seurat, and the data were scaled using “ScaleData” to perform dimensional reduction. First, PCA was applied, and a representative number of relevant dimensions was selected to construct clusters that identify each cell type. Both RNA and ADT profiles for all groups were integrated using the “IntegrateData” function. The data was again normalized, the variable features were recalculated and scaled, and the clusters were identified for the integrated data. Cell identification was performed manually based on the ADT content and subsequently classified according to the RNA content. Differential expression was calculated using Seurat’s “FindMarkers” function with the Wilcoxon rank-sum test for both RNA and ADT data. Re-clustering of some groups was performed to obtain more detailed information.

## Acknowledgements

This grant was supported by NIAID’s grants R01AI175567, R01AI169460, R01AI169460-02S1, R21AI156490, R21AI153920, and R56AI110457, NCI’s grant, and NIH Office of the Director’s grant S10 OD02512. We are also thankful to Thomas Jefferson University’s “Flow Cytometry and Human Monitoring”, “Cancer Genomics”, and “Laboratory Animal Services” Shared Resources and to the University of Pennsylvania’s “Penn Cytomics and Cell Sorting Resource Laboratory” and their personnel for their support.

## Author contributions

C.R.M-S. conceived the study, developed the study methodology, designed and made the genetically modified mice by iGONAD, performed most of the experiments, analyzed the data, supervised scSeq analysis, and wrote the manuscript. L.J.S. conceived the study, developed the study methodology, directed and supervised the overall project, wrote the manuscript, and obtained funding. M.I.R. performed all the scSeq bioinformatics. N.H. and M.G. performed some experiments as part of their master’s thesis under C.R.M-S. and L.J.S.’s supervision. L.T. assisted in generating the genetically modified mice generated by iGONAD and produced ECTV stocks. L.T. and O.T. were responsible for breeding and genotyping all the mouse strains. L.T., M.K., J. C., O. T., S. K., and E. T-G. helped with or performed some experiments. H.R. prepared West Nile virus stocks and provided insights about the model. R.A. provided ideas and insights.

