## Supplemental figures and material for "Interferon-α and -β subtypes have temporally distinct roles in containing viral spread and protecting vital organs"

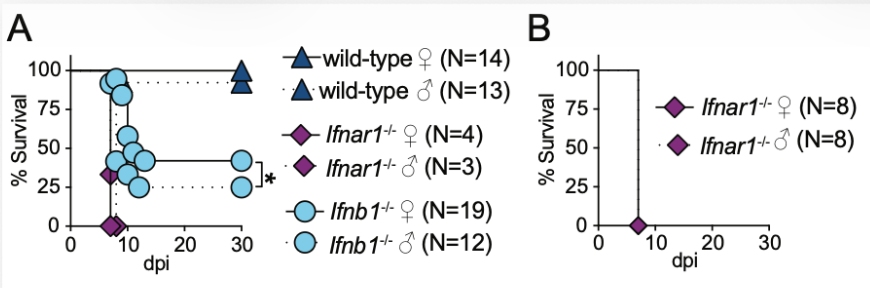

**Supplemental Figure 1. IFN-I signaling partially contributes to the male-associated increase in susceptibility to mousepox. A-B**) Survival of the indicated mice infected in the footpad with 3,000 pfu of ECTV. The data correspond to four (**A**) or two (**B**) independent experiments combined. P values are compared to wild-type mice or the indicated groups by Log-rank (Mantel-Cox) analysis.

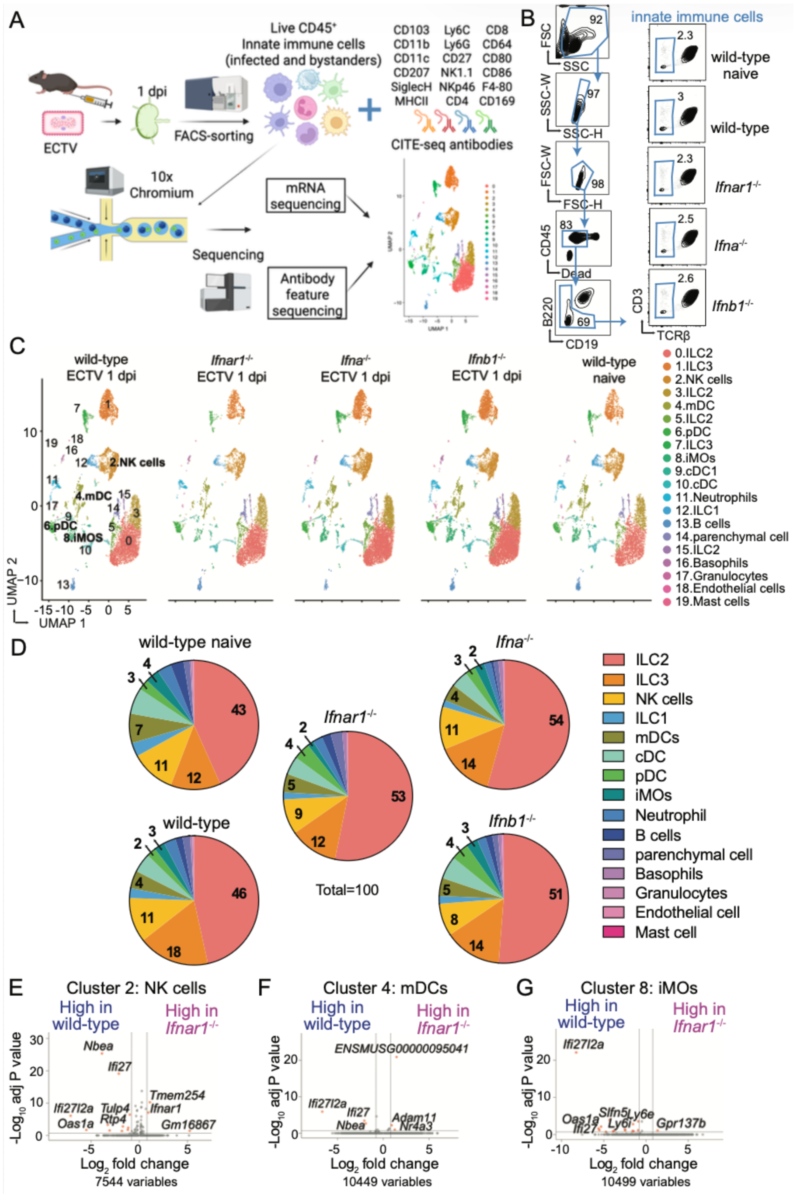

**Supplemental Figure 2. IFN-I partially primes innate immune cells in the dLN at one dpi with ECTV. A**) CITEseq experimental layout (created with BioRender). **B**) FACS plots showing the gating strategy for the sorting of live innate immune cells (CD45$^+$TCRβ$^-$CD3$^-$B220$^-$CD19$^-$) from pooled draining lymph nodes at one dpi from the indicated mice infected with 3,000 pfu of ECTV in both rear footpads or from naïve wild-type controls. **C**) UMAP clustering and cell annotation based on cell-specific protein markers and transcript expression. **D**) Percentages of the indicated cell populations of total sequenced cells. **E-G**) Volcano plots showing differentially expressed genes of clusters annotated as NK cells (**E**, NK1.1$^+$ NKp46$^+$*Eomes*$^+$SiglecH$^-$MHCII$^-$Ly6G$^-$CD103$^-$), mDCs (**F**, MHCII$^{high}$CD11c$^+$ CD80$^+$ CD86$^+$ CD207$^+$SiglecH$^-$Ly6C$^-$CD11b$^-$Ly6G$^-$NK1.1$^-$CD103$^-$) or iMOs (**G**, Ly6C$^+$CD11b$^+$CD11c$^+$CD80$^+$SiglecH$^-$ Ly6G$^-$NK1.1$^-$CD103$^-$) from infected wild-type mice compared to those from infected *Ifnar1*$^{-/-}$ mice (red dots: Log2 fold change=1, adj P<0.2, gray dots: unchanged).

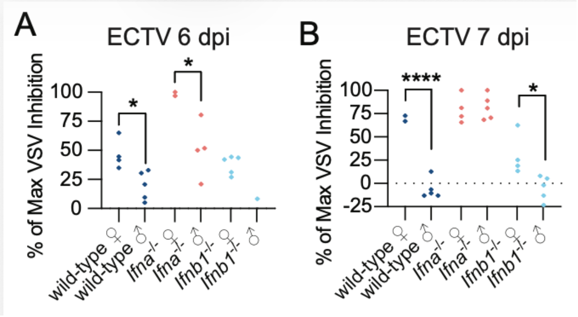

**Supplemental Figure 3. Systemic IFN-I signaling is delayed or reduced in male compared to female mice infected with ECTV or WNV. A-B**) IFN-I bioactivity in the sera obtained from the indicated mice infected with ECTV at six (**A**) or seven dpi (**B**) was quantified by inhibition of VSV replication in mouse L929 cells. The data correspond to two independent experiments combined for **A-B**. Each symbol depicts an individual mouse, and P values are compared to the indicated groups by t-test with Welch's correction.

Box1. Material and equipment.

| REAGENT or RESOURCE | SOURCE | IDENTIFIER |
|---|---|---|
| Antibodies | | |
| Mouse monoclonal antibody anti-mouse anti-interferon beta, clone HDβ-4A7 | Leinco PMC4444312 | I-1182 |
| Armenian hamster monoclonal antibody anti-mouse anti-interferon alpha, clone TIF-3C5 | Leinco PMC4444312 | I-1183 |
| Mouse monoclonal antibody Buv395 anti-mouse CD45.2, clone 104 | BioLegend | Cat# 564616, RRID: AB_893350 |
| Rat monoclonal antibody FITC anti-mouse CD3, clone 17A2 | BioLegend | Cat# 100203, RRID: AB_312660 |
| Armenian hamster monoclonal antibody APC anti-mouse TCRβ, clone H57-597 | BioLegend | Cat# 109211, RRID: AB_313434 |
| Rat monoclonal antibody PE anti-mouse B220, clone RA3-6B2 | BioLegend | Cat# 103207, RRID: AB_312992 |
| Rat monoclonal antibody BV605 anti-mouse CD19, clone 6D5 | BioLegend | Cat# 115540, RRID: AB_2563067 |
| Rat monoclonal antibody PE anti-mouse Ly6C, clone HK1.4 | BioLegend | Cat# 128008, RRID:AB_1186132 |
| Rat monoclonal antibody BUV395 anti-mouse CD11b, clone M1/70 | BD Biosciences | Cat# 565976, RRID:AB_2721166 |
| Armenian hamster monoclonal antibody PE-Cy7 anti-mouse CD11c, clone N418 | BioLegend | Cat# 117318 RRID: AB_493569 |
| Rat monoclonal antibody PerCP/Cy5.5 anti-mouse I-A/I-E, clone M5/114.15.2 | BioLegend | Cat# 107626, RRID:AB_2191071 |
| Rat monoclonal antibody Pacific Blue anti-mouse Ly6C/Ly6G (Gr-1), clone RB6-8C5 | BioLegend | Cat# 108430, RRID:AB_893556 |
| Rat monoclonal antibody BV785 anti-mouse CD19, clone 6D5 | BioLegend | Cat# 115543, RRID:AB_11218994 |
| Rat monoclonal antibody anti-mouse CD16/CD32, clone 93 | BioLegend | Cat#101302 |
| Armenian hamster monoclonal antibody BV605 anti-mouse TCRβ, clone H57-597 | BioLegend | Cat# 109241, RRID:AB_2629563 |
| Mouse monoclonal antibody BV605 anti-mouse NK 1.1, clone PK136 | BioLegend | Cat# 108753, RRID: AB_2686977 |
| Armenian Hamster monoclonal antibody TTCATTAGCCCGCTG-barcoded anti-mouse CD103, clone 2E7 | BioLegend | Cat# 121445, RRID: AB_2876437 |
| Rat monoclonal antibody TGAAGGCTCATTTGT-barcoded anti-mouse CD11b, clone M1/70 | BioLegend | Cat# 101273, RRID: AB_2819781 |
| Armenian Hamster monoclonal antibody GTTATGGACGCTTGC-barcoded anti-mouse CD11c, clone N418 | BioLegend | Cat# 117359, RRID: AB_2813987 |
| Mouse monoclonal antibody CGATTTGTATTCCCT-barcoded anti-mouse CD207, clone 4C7 | BioLegend | Cat# 144213, RRID: AB_2876492 |
| Armenian Hamster monoclonal antibody CAAGGTATGTCACTG-barcoded anti-mouse CD27, clone LG.3A10 | BioLegend | Cat# 124247, RRID: AB_2832449 |
| Mouse monoclonal antibody AGCAATTAACGGGAG-barcoded anti-mouse CD64, clone X54-5/7.1 | BioLegend | Cat# 139329, RRID: AB_2832505 |
| Armenian Hamster monoclonal antibody GACCCGGTGTCATTT-barcoded anti-mouse CD80, clone 16-10A1 | BioLegend | Cat# 104755, RRID: AB_2819807 |

| | | |
|---|---|---|
| Rat monoclonal antibody CTGGATTTGTGTATC-barcoded anti-mouse CD86, clone GL-1 | BioLegend | Cat# 105057, RRID: AB_2832345 |
| Rat monoclonal antibody TTAACTTCAGCCCGT-barcoded anti-mouse F4-80, clone BM8 | BioLegend | Cat# 123155, RRID: AB_2819847 |
| Rat monoclonal antibody AAGTCGTGAGGCATG-barcoded anti-mouse Ly6C, clone HK1.4 | BioLegend | Cat# 128053, RRID: AB_2832462 |
| Rat monoclonal antibody ACATTGACGCAACTA-barcoded anti-mouse Ly6G, clone 1A8 | BioLegend | Cat# 127659, RRID: AB_2819864 |
| Rat monoclonal antibody GGTCACCAGTATGAT-barcoded anti-mouse I-A/I-E, clone M5/114.15.2 | BioLegend | Cat# 107657, RRID: AB_2832367 |
| Mouse monoclonal antibody GTAACATTACTCGTC-barcoded anti-mouse NK1.1, clone PK136 | BioLegend | Cat# 108763, RRID: AB_2819814 |
| Rat monoclonal antibody CCCTTTCACCTCGAA-barcoded anti-mouse NKp46, clone 29A1.4 | BioLegend | Cat# 137641, RRID: AB_2860685 |
| Rat monoclonal antibody CCGCACCTACATTAG-barcoded anti-mouse SiglecH, clone 551 | BioLegend | Cat# 129619, RRID: AB_2922464 |
| Rat monoclonal antibody ATTGACGACAGTCAT-barcoded anti-mouse CD169, clone 3D6.112 | BioLegend | Cat# 142429, RRID: AB_2876485 |
| Rat monoclonal antibody TACCCGTAATAGCGT-barcoded anti-mouse CD8a, clone 53-6.7 | BioLegend | Cat# 100783, RRID: AB_2832269 |
| Rat monoclonal antibody AACAAGACCCTTGAG-barcoded anti-mouse CD4, clone RM4-5 | BioLegend | Cat# 100573, RRID: AB_2813914 |
| Rat monoclonal antibody AAGTCAGGTTCGTTT-barcoded κ isotype control, clone RTK2758 | BioLegend | Cat# 400581, RRID: AB_3097104 |
| Virus Strains | | |
| ECTV Moscow | ATCC | VR-1374 |
| ECTV ΔEVM166 | PMC2292233 | |
| ECTV-GFP | PMC2233669 | |
| Chemicals, Peptides, and Recombinant Proteins | | |
| Zombie Violet Fixable Viability kit | BioLegend | Cat# 423113 |
| Mouse interferon alpha A | PBL Assay Science | Cat# 12100 |
| Mouse interferon alpha 1 | PBL Assay Science | Cat# 12105 |
| Mouse interferon beta | PBL Assay Science | Cat# 12400 |
| DMEM media | CORNING | 10-013-CV |
| RPMI media | CORNING | 10-040-CV |
| Penicillin Streptomycin Solution, 100x | CORNING | 30-002-Cl |
| GlutaMAX 100x | Gibco | 35050-061 |
| Hepes 1M | CORNING | 25-060-Cl |
| Liberase™ | Roche | 05 401 119 001 |
| Trizol | Invitrogen | Cat# 15596018 |
| Critical Commercial Assays | | |
| cDNA synthesis with High Capacity cDNA Reverse Transcription Kit | Applied Biosystems™ Thermo Fisher Scientific | 4368814 |
| iTaq™ Universal SYBR® Green PCR Supermix | BioRad | 172-5124 |
| RNA Clean and Concentrator™-5, DNaseI | Zymo Research | R1014 |
| Dual Index Kit NT Set A, 96 rxn | 10x Genomics | 1000242 |
| Chromium Next GEM Chip G Single Cell Kit, 16 rxns | 10x Genomics | 1000127 |
| Chromium Next GEM Single Cell 3' Kit v3.1, 16 rxns | 10x Genomics | PN-1000268 |
| 3' Feature Barcode Kit, 16 rxns | 10x Genomics | PN-1000262 |

| | | |
|---|---|---|
| Experimental Models: Cell Lines | | |
| Monkey *C. aethiops* epithelial kidney BS-C-1 cells | ATCC | CCL-26 |
| Mouse *M. musculus* subcutaneous connective tissue L929 | ATCC | CCL-1 |
| Hamster *M. auratus* fibroblast BHK-21 | ATCC | CCL-10 |
| Mosquito *A. albopictus* larva C6/36 | ATCC | CRL-1660 |
| Mouse *M. musculus* hybridoma A9 | ATCC | CRL-1811 |
| Monkey *C. aethiops* kidney epithelial Vero | ATCC | CCL-81 |
| Experimental Models: Organisms/Strains | | |
| Mouse: C57BL/6NCrl | Charles River | 027 |
| Mouse: C57BL/6 *Ifnar1*$^{-/-}$ mice | Thomas Moran (Mount Sinai School of Medicine, New York, NY) | |
| Mouse: C57BL/6;129P2-Irf7tm1Ttg/TtgRbrc - *Irf7*$^{-/-}$ | Dr. Tadatsugu Taniguchi, PMID: 15800576 | |
| Mouse: C57BL/6.C-Tg(CMV-cre)1Cgn/J - CMV-Cre | Jackson Laboratory | JAX: 006054 |
| Mouse: C57BL/6 *Ifnb1*$^{-/-}$ | PMC10177031 | |
| Mouse: C57BL/6 *Ifna4,b1*$^{-/-}$ | PMC10177031 | |
| Mouse: C57BL/6 *Ifna4*$^{-/-}$ | PMC10177031 | |
| Mouse: C57BL/6 *Ifna*$^{\Delta 3}$ | PMC10177031 | |
| Mouse: C57BL/6 *Ifna*$^{\Delta 6}$ | PMC10177031 | |
| Mouse: C57BL/6 *Ifna*$^{-/-}$ | Present paper | Offspring from crossing of C57BL/6 CMV-Cre and C57BL/6 *Ifna*$^{fl/fl}$ |
| Mouse: C57BL/6 *Ifna*$^{fl/fl}$ | PMC10177031 | |
| Mouse: C57BL/6 *Ifnb1*$^{\Delta PRDII}$ | PMC10177031 | |
| Mouse: C57BL/6 *Ifnb1*$^{\Delta PRDII}$*Irf7*$^{-/-}$ | Present paper | Offspring from crossing of C57BL/6 *Ifnb1*$^{\Delta PRDII}$ and C57BL/6 *Irf7*$^{-/-}$ |
| Oligonucleotides | | |
| Primer *Gapdh* forward | IDT | tgtccgtcgtggatctgac |
| Primer *Gapdh* reverse | IDT | cctgcttcaccaccttcttg |
| Primer *Ifnb1* forward | IDT | ctggcttccatcatgaacaa |
| Primer *Ifnb1* reverse | IDT | agagggctgtggtggagaa |
| Primer *Ifna4* forward | IDT | tcaagccatccttgtgctaa |
| Primer *Ifna4* reverse | IDT | gtcttttgatgtgaagaggttcaa |
| Primer *Ifna* non-a4 forward (II *Ifna*) | IDT | ARSYtgtStgatgcaRcagg t |
| Primer *Ifna* non-a4 reverse (II *Ifna*) | IDT | ggWacacagtgatcctgtgg |
| Software and Algorithms | | |
| Prism | GraphPad Software | version 10 |
| cellranger | 10x Genomics | version 8.0.1 |
| Seurat package | PMC4430369 | version 5.2.0 |
| RStudio | | version 4.5.5.1 |
| FlowJo™ | Treestar | version 10 |
| Other | | |
| TissueLyzer II | Qiagen | 85300 |
| Thermocycler CFX96 Real-Time System | BioRad | |

| | |
|---|---|
| Chromium X | 10x Genomics |
